# Cell-to-cell transmission of *Brucella abortus*

**DOI:** 10.1101/2025.10.15.682732

**Authors:** José F. Campos-Godínez, Carlos Chacón-Díaz, Edgardo Moreno, Esteban Chaves-Olarte, Pamela Altamirano-Silva

## Abstract

*Brucella abortus* is a persistent intracellular pathogen capable of evading early immune responses. To explore the mechanism of exit of this bacterium, we monitored *Brucella* infection by live cell imaging for up to 100 hours. This approach uncovered various modes of bacterial spreading, including individual organisms, bacteria-containing vacuoles, and direct transmission between cells (called cell-to-cell transmission), each fueling new infections. This dissemination occurred regardless of antibiotic presence in the extracellular milieu. While receptor blockade using heparin suppressed free bacterial infections, it promoted cell-to-cell transmission, dependent on cell density. Cells participating in intercellular transmission survived longer than their non-infected counterparts. These observations underscore the adaptability of *Brucella* for propagating across diverse conditions.

**Importance:** The intracellular life cycle of pathogenic bacteria encompasses adhesion, invasion, intracellular replication, and eventual exit from host cells. The exit phase is a critical step for pathogen persistence and dissemination to new hosts. Despite its relevance, the egress mechanisms of *Brucella* spp. remain largely unexplored. In this study, we describe three distinct modes of *Brucella abortus* exit: individual bacteria, bacterial clumps, and through direct cell-to-cell transmission. The latter may facilitate immune evasion by bypassing extracellular exposure, thereby contributing to the stealthy survival strategy of *Brucella*. The presence of multiple egress mechanisms suggests that *B. abortus* tailors its exit strategies according to its intracellular biogenesis and environmental cues.These findings not only advance our understanding of *Brucella* pathogenesis but also highlight potential avenues for targeting bacterial dissemination and persistence.

## Introduction

*Brucella abortus* invades, persists, and replicates within host cells, trafficking through vacuolar compartments and replicating in endoplasmic reticulum-derived structures (1-3). At later infection stages, host cells activate Rho GTPases and reorganize actin, releasing individual bacteria or *Brucella-*containing vacuole-derived extracellular clusters (BCVEC) from autophagic vacuoles. BCVEC, containing 10–250 bacteria, are enriched with Lamp-1 and actin and show high infectivity. They are either exposed or enclosed in host-derived membranes (4). Beyond releasing free bacteria and BCVECs, we show that infected cells facilitate bacterial transmission via intercellular contacts, a phenomenon not previously documented in Brucella pathogenesis.

## Results

Gentamicin has minimal impact on *B. abortus* replication in HeLa cells, causing only a slight reduction in bacterial egress **(Fig. 1A)**. In addition to protrusion of single bacteria and, heavily infected cells promote transmission through cell-to-cell contact in approximately 25% of infected HeLa cells **(Fig. 2A and B)**. This transmission emerges around 60 h. post-infection (p.i.) and remains unaffected by gentamicin **(Fig. 2B)**. Recipient cells become fully loaded over 44.2 ± 4.6 h. after transfer begins (n = 30). Transmission occurs from one donor to a single recipient in approximately 72% of cases, while 28% involve two distinct recipients **(Fig. 3A and B)**. Transmission to more than two cells is rare. After four days, propidium iodide staining shows that approximately 74% of transmitting cells retain membrane integrity, compared to 78% of non-infected cells that exhibit permeability **(Fig. 3C)**. Cell-to-cell transmission requires physical contact, as infections drop from approximately 37% under 80–95% confluency to none under 40–70%, while alternative transmission routes increase **(Fig. 3D)**. Heparin, a structural analogue of heparan sulfate proteoglycans involved in *B. abortus* adhesion (5), blocks receptors in naïve cells, reducing individual reinfection by approximately 15%, enhancing cell-to-cell transmission by 16%, and leaving BCVEC spread unaffected, all independent of gentamicin **(Fig. 3E and F)**.

**Figure 1.**
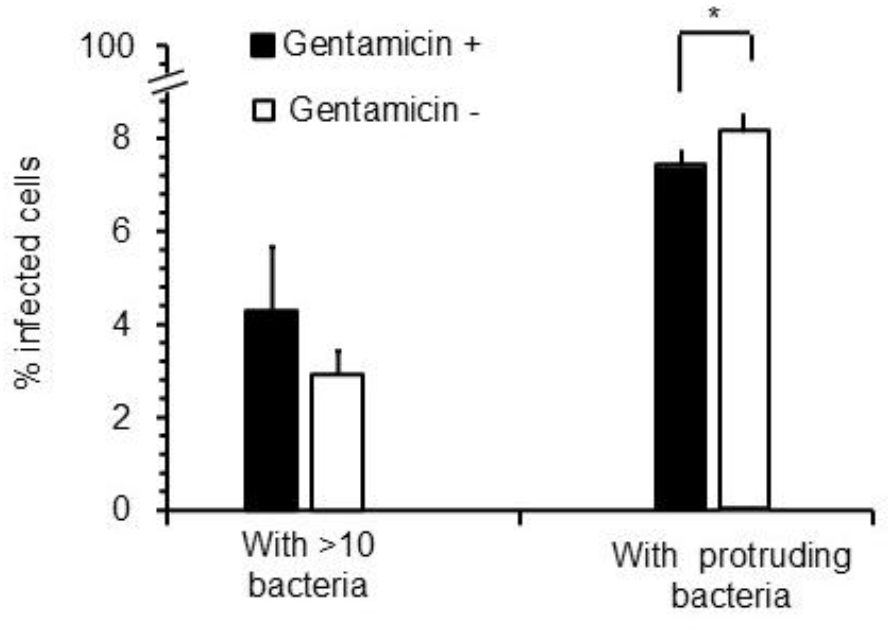
*B. abortus* replication and egress. A) HeLa cells were infected with *B*.*abortus* RFP and incubated with gentamicin and at 48 h p.i. infected cells were incubated with or without gentamicin. Infected cells or cells with protruding bacteria were recorded and counted.

**Figure 2.**
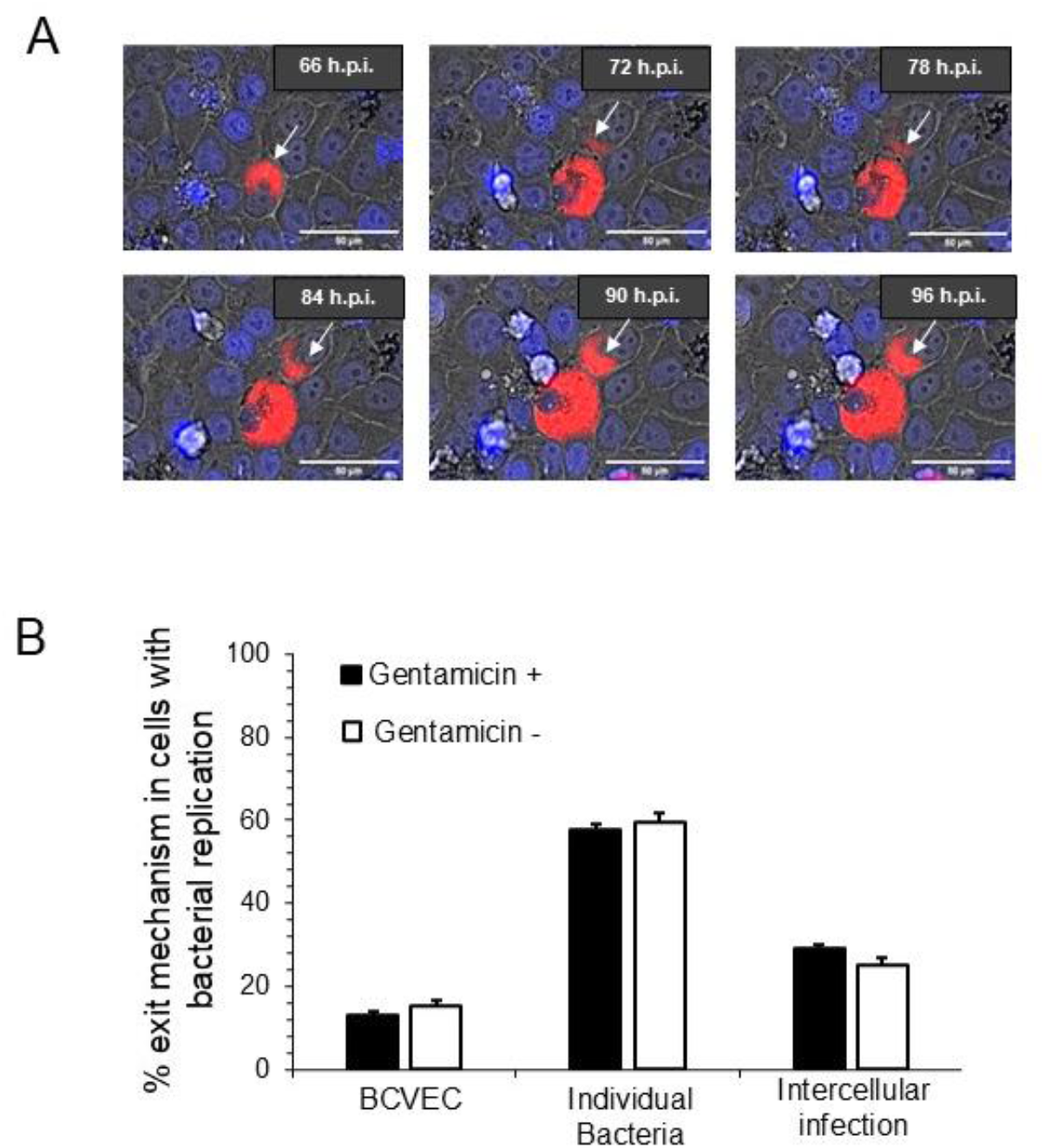
*B. abortus* cell-to-cell transmission. A) Infection spreads through HeLa cell contacts. Arrows indicate transmission sites and subsequent replication in recipient cells. All frames show the same field at different times. B) BCVEC and single bacteria initiate new or cell-to-cell infections with or without gentamicin. Infection rates show no significant difference between treated and untreated conditions.

**Figure 3.**
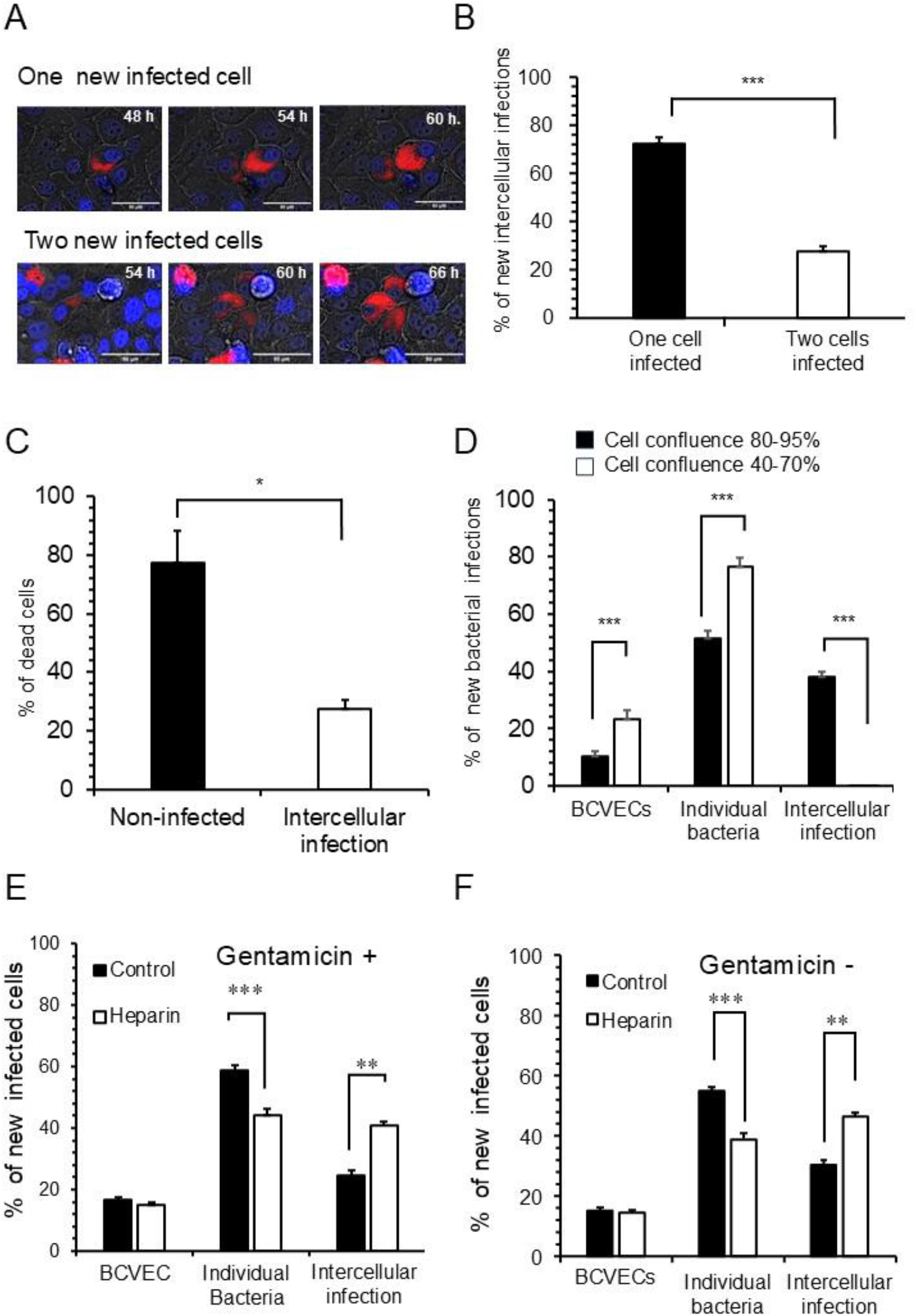
*B. abortus* cell-to-cell transmission prolongs host cell survival. A single infected HeLa cell initiated one or two transmission events. A) Quantified in panel “B”. C) Infected donor cells showed greater survival than non-infected cells. D) At 48 h. post-infection, cells were dissociated and used to infect non-infected cultures at different confluence levels. BCVECs, individual bacteria, and intercellular transmission were tracked by real-time fluorescence microscopy until 100 h. E) Newly infected cells were distributed among BCVECs, protruding bacteria, or cell-to-cell transmission under heparin and gentamicin treatment. F) A parallel experiment performed without gentamicin (F).

## Discussion

*Brucella* infection mechanisms are critical because they enable the bacteria to evade immune defenses. In addition to single bacteria and BCVEC, here we showed that *Brucella* also propagates through cell-to-cell contacts **(Figure 4)**. Like other infected cells, donor cells live longer than their non-infected counterparts (6,7), demonstrating that intracellular brucellae influence host cell cycle through a still unresolved mechanism. Macrophage phagocytosis of infected neutrophils, individual bacteria, or bacterial clusters serves a distinct function for spreading *Brucella* (4,8). Whether the transmission mode influences intracellular bacterial trafficking remains unknown. Several pathogens, including *Brucella*, rely on heparan-sulfate proteoglycans to adhere to host cells, facilitating the onset and persistence of infection (9,10). Our findings confirm that heparin disrupts individual bacterial adhesion as a key infection step. Therefore, alternative receptors may participate in BCVEC transmission. Likewise, suppressing adhesion with heparin increases cell-to-cell transmission, indicating that *B. abortus* adapts its exit strategies to its intracellular biogenesis.

**Figure 4.**
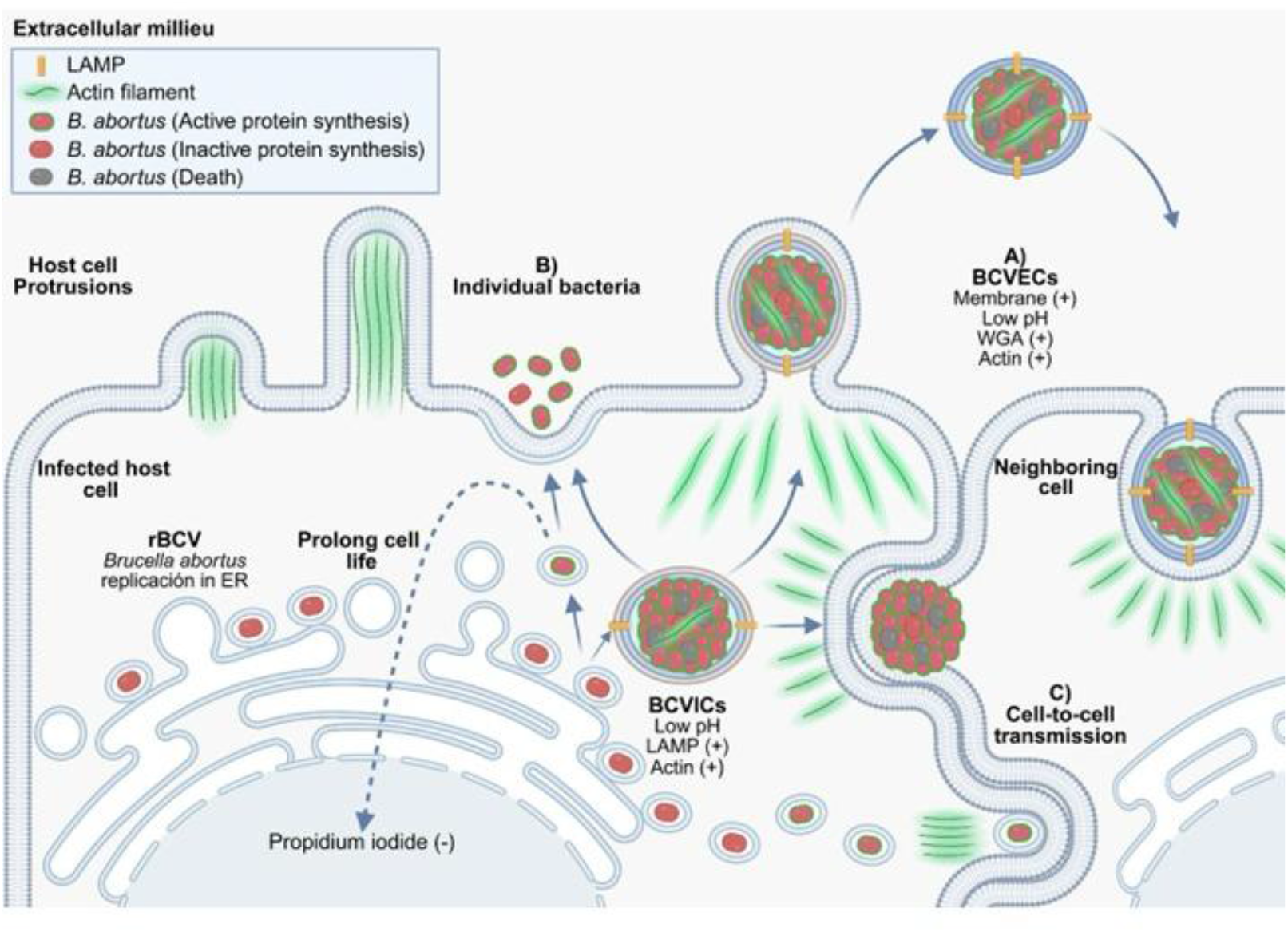
Model showing *B. abortus* egress from infected cells. After extensive replication at the endoplasmic reticulum-derived compartments, *B. abortus* organisms cluster in an acidic vacuole within characteristics of autophagosomes (BCVIC) as proposed by Starr et al (Starr et al., 2012). During this period, the replicating bacteria prolongs the life of the infected cell. Then, *B. abortus* egress from the host cell by three mechanisms: (A) released BCVEC, (B) individual bacteria and (C) cell-to-cell transmission. Figure modified from Campos-Godinez et al, 2025 (Under submission).

## Materials and Methods

HeLa cells infected with *B. abortus* 2308W-RFP were monitored by real-time fluorescence microscopy until 100 h. p.i. (4). At various times, cells were untreated or treated with 5 μg/mL gentamicin or 25 μg/mL heparin sodium salt (SIGMA) from 48–100 h. p.i.. At 48 h., infected cells were dissociated with 0.05% Trypsin-EDTA (Gibco) for 10 minutes to enable cell-to-cell transmission, then resuspended in DMEM with 5 μg/mL gentamicin and seeded onto wells with non-infected cells at 40–70% or 80–95% confluence. Live-cell imaging was performed from 48 to 100 h. using a BioTek Lionheart FX Automated Microscope at 37 °C under 5% CO_2_, with images acquired every 4 h. Image contrast was uniformly increased by 15% using Adobe Photoshop® Hue adjustment. Data were analyzed with GraphPad Prism 8.0.1, and statistical significance was determined by one-way ANOVA followed by Tukey’s post-hoc test (11).

## References

1. Moreno, E., and Moriyon, I. 2006. The Genus Brucella In The Prokaryotes, M. Dworkin, S. Falcow, E. Rosenberg, K.-H. Schleifer, and E. Stackebrandt, eds. (Springer, New York), pp. 315–456.

2. Pizarro-Cerdá, J., Méresse, S., Parton, R.G., van der Goot, G., Sola-Landa, A., Lopez-Goñi, I., Moreno, E., and Gorvel, J.P. 1998. Brucella abortus transits through the autophagic pathway and replicates in the endoplasmic reticulum of nonprofessional phagocytes. Infect Immun 66, 5711–5724.

3. Celli, J. 2015. The changing nature of the Brucella-containing vacuole. Cell Microbiol 17, 951–958. 10.1111/cmi.12452.

4. F. Campos-Godínez, A. Hernández-Saborío, T. Vargas-Moya, V. Gómez-Vargas, D. Espinoza-Villagra, I. Sandoval, M.-P. Rojas-Salas, M. López-Hernández, R. Vega-Arce, R. Pereira-Reyes, M. Prado, C. Chacón-Díaz, E. Moreno, E. Chaves-Olarte, P. Altamirano-Silva. Brucella abortus egresses from host cells in infective clumps through an actin-dependent mechanism. Under submission. mBio. 2025.

5. Shriver, Z., Capila, I., Venkataraman, G., and Sasisekharan, R. 2012. Heparin and heparan sulfate: analyzing structure and microheterogeneity. Handb Exp Pharmacol, 159–176. 10.1007/978-3-642-23056-1_8.

6. Chaves-Olarte, E., Guzmán-Verri, C., Méresse, S., Desjardins, M., Pizarro-Cerdá, J., Badilla, J., Gorvel, J.P., and Moreno, E. 2002. Activation of Rho and Rab GTPases dissociates Brucella abortus internalization from intracellular trafficking. Cell Microbiol 4, 663–676.

7. Barquero-Calvo, E., Chaves-Olarte, E., Weiss, D.S., Guzmán-Verri, C., Chacón-Díaz, C., Rucavado, A., Moriyón, I., and Moreno, E. 2007. Brucella abortus uses a stealthy strategy to avoid activation of the innate immune system during the onset of infection. PLoS One 2, e631. 10.1371/journal.pone.0000631.

8. Moreno, E., and Barquero-Calvo, E. 2020. The Role of Neutrophils in Brucellosis. Microbiol Mol Biol Rev 84. 10.1128/MMBR.00048-20.

9. Castañeda-Roldán, E.I., Avelino-Flores, F., Dall’Agnol, M., Freer, E., Cedillo, L., Dornand, J., and Girón, J.A. 2004. Adherence of Brucella to human epithelial cells and macrophages is mediated by sialic acid residues. Cell Microbiol 6, 435–445. 10.1111/j.1462-5822.2004.00372.x.

10. del C Rocha-Gracia, R., Castañeda-Roldán, E.I., Giono-Cerezo, S., and Girón, J.A. 2002. Brucella sp. bind to sialic acid residues on human and animal red blood cells. FEMS Microbiol Lett 213, 219–224. 10.1111/j.1574-6968.2002.tb11309.x.

11. McHugh, M.L. 2011. Multiple comparison analysis testing in ANOVA. Biochem Med (Zagreb) 21, 203–209. 10.11613/bm.2011.029.

